# Innovative Nitinol Lumbar Vertebral Implants: Enhancing Biocompatibility and Mechanical Properties through Advanced Braiding Techniques

**DOI:** 10.1101/2025.02.25.640102

**Authors:** Mahdis Parsafar, Alireza Tehranian, Ramin Ardalani

**Affiliations:** Department of Biomedical Engineering, Faculty of Paramedical Sciences, Tehran Medical Sciences, Islamic Azad University, Tehran, Iran

**Keywords:** Vertebral implant, Corrosion, Nitinol, Biocompatibility

## Abstract

The increasing occurrence of orthopedic disorders calls for advancements in techniques and implant technologies. The sensitive nature of the spinal cord highlights the need for implants with specific mechanical and biological properties that imitate those of bone. Nitinol alloy, known for its super-elasticity, bone-like elastic modulus, and excellent compatibility with living tissues is emerging as a favored material for orthopedic implants. In this research, a nitinol lumbar vertebral implant was created using a two-step procedure involving rod fabrication through sintering and braiding techniques focusing on assessing the effects of incorporating braiding into the structure. This method aimed to achieve both mechanical and biological properties simultaneously. In previous techniques, there was a greater focus on one of these properties. However, in this approach, the addition of the braid was an innovative way to append and enhance the biological properties. Examination under Scanning Electron Microscopy revealed a pore size of 310 micrometers in the samples while Atomic Force Microscopy showed a roughness of 971.1 nm both properties are proper for bone cells. Corrosion resistance was evaluated using Electrochemical Impedance Spectroscopy and the measured amount was 7366 ohm.cm2. Biological tests confirmed that the samples were biocompatible and non-toxic, with a cell viability rate of 95.1% observed over a period of 21 days. The development of this type of implant aims to improve the treatment outcomes for vertebral area surgeries.

## 1. Introduction

The human spine, renowned for its structural robustness, plays a critical role in protecting the spinal cord and nerve roots [1]. Its favorable mechanical qualities and high fracture toughness enable it to facilitate motion and safeguard delicate internal organs [2]. Among the spinal regions, the lumbar area, which houses various components such as visceral organs, is a significant source of low back pain (LBP) [3]. LBP is characterized by discomfort originating from the buttock region and traveling down the trajectory of the sciatic nerve [4, 5]. Over the past twenty-five years, there has been a notable rise in the occurrence of LBP, with the worldwide incidence recorded at 568.4 million cases [6, 7]. Treating and managing back pain poses considerable financial and medical challenges due to the delicate and vulnerable nature of the spinal cord [8].

In response to these challenges, significant research over the last two decades has focused on evaluating different bone replacement materials and providing bone substitutes. These materials often include bioactive ceramics and biological or synthetic polymers. However, fabricating orthopedic implants with metal alloys that exhibit exceptional mechanical properties has proven more practical [9, 10, 11, 12]. Spinal implants must possess two crucial attributes: mechanical qualities, including bio-functionality, and biological traits, including biocompatibility. Among various alloys, Nitinol has been selected for orthopedic implants due to its ability to undergo a phase change called the martensite-austenite transformation, affecting the implant’s mechanical properties to closely mimic those of bone. This transition is often prompted by stimuli linked to temperature and/or stress [13].

Nitinol refers to a group of intermetallic materials mostly composed of titanium and nickel in about equal proportions. Its shape memory and superelastic capabilities are attributed to its thermoelastic martensitic transition [14, 15, 16]. Nitinol’s diverse beneficial features make it a highly promising material for biomedical applications.

Conversely, due to the invasiveness of surgery, a promising method for bone regeneration is bone tissue engineering (BTE). Synthetic extracellular matrix (ECM) and osteogenic activity stimulate bone regeneration [17]. Ravindran et al. have shown that ECM ranks as the fourth most significant element in the development of BTE [18]. Consequently, the induction of osteogenic activity has been demonstrated to enhance the process of bone regeneration [19, 20]. A porous structure within these materials allows infiltration into host tissue, stimulating cellular proliferation, capillary development, and bone tissue creation [21, 22, 23].

Developing NiTi implants for lumbar vertebrae is a substantial challenge in orthopedic biomedical engineering, involving further advancements. Nitinol is the preferred material for this application due to its superior performance features and required attributes. The current study employed sintering and braiding techniques simultaneously to fabricate a Nitinol implant in the shape of a lumbar vertebra. Constructing a framework that possesses both flexibility and endurance necessitates the intricate interlacing of several strands of Nitinol wire. This novel braiding method is distinguished by its simplicity and cost-effectiveness, rendering it an innovative and pragmatic approach to production. This methodology enables precise manipulation of the implant’s dimensions and configuration, making it suitable for the fabrication of customized implants with improved mechanical and biological characteristics. It is important to note that using the rod without braid (WOB) would increase the risk of shielding stress effects, reducing biocompatibility and adhesion, which would diminish their overall benefits [24]. However, the absence of a rod would lead to unfavorable mechanical corrosion outcomes. A two-step approach was used, involving the initial fabrication of a rod, followed by the secondary procedure of wire encircling the rod.

The main aim of this work is to enhance the orthopedic biomedical engineering field through a meticulous investigation of the potential suitability of Nitinol as a material for vertebral implants. This research seeks to augment orthopedic technology by thoroughly examining the potential applications of Nitinol as a material for implants. Additionally, the present work introduces a novel production concept for Nitinol, intending to facilitate more comprehensive investigations in this domain.

## 2. Material and methods

### 2.1. Preparation process

The elements required to achieve a titanium concentration of 50.0% with nickel. It is noteworthy to state that a porosity maker was included to augment the porosity level to 50.0%, As shown in Table 1. Nitinol wire from the Baoji Hanz company was employed for this experiment, and its diameter was 0.25 mm. Urea’s size showed a wide range of variability, from 100 to 600 μm. 150 MPa of pressure was imposed on the mixture. The specimen spent two hours undergoing thermal treatment in an argon-filled furnace at a temperature of 1050°C. Urea will be eliminated through evaporation, whereas titanium hydride will undergo decomposition, when the temperature reaches 200 and 500°C, respectively. It is important to mention that the material was exposed to both temperatures for one hour. The resulting sample measures 7 mm in height and 5 mm in diameter without the braid (WOB). In the next step, for obtaining samples with braid (WB), employing a machine for the braiding process proved to be immensely efficient in terms of time. The dimensions of the specimen were used to create a pattern, which was then woven together and placed in a cylinder around its circumference. Figures 1 and 2 show images of the finalized examples. A pair of wires were used to produce this texture. The rod (WOB) serves as the primary focus of the structure. As can be seen in Figure 1, the first wire makes its way down from the green area. After that, it leaves the green zone, with the “+” sign denoting the outside and the “-” symbol on the inside. This texture was produced by using two wires. The rod makes up the design’s focal point. As shown in Figure 1, the first cable enters the underground zone in the green area, then travels via green area 2, and so on until it emerges in green area 5. Next, the wire goes through a similar process inside the blue areas, but in reverse. Red lines on the rod indicate the exact points where two wires cross each other. Figure 2 shows the braided pattern without the rod. It’s crucial to remember that this method was developed to improve mechanical and biological behavior. The presence of a braided cover helps biological properties and minimizing the use of Nitinol wire to the greatest extent possible proved to be a cost-effective approach while maintaining optimal performance. The following sections will provide a comprehensive account of the procedures involved in the development and production of the braid. Collaboration is advantageous and valuable in terms of fostering cooperation.

**Figure 1.**
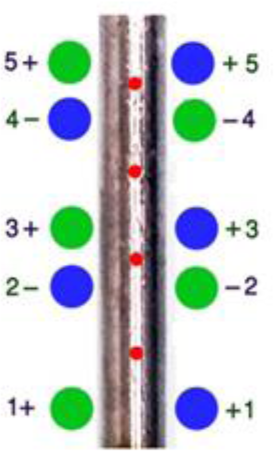
The place where the wires pass the process Through the green and blue circles around the cylinder.

**Figure 2.**
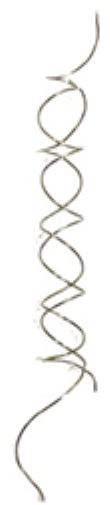
The design of the cover made after of Figure 1.

**Table 1.**
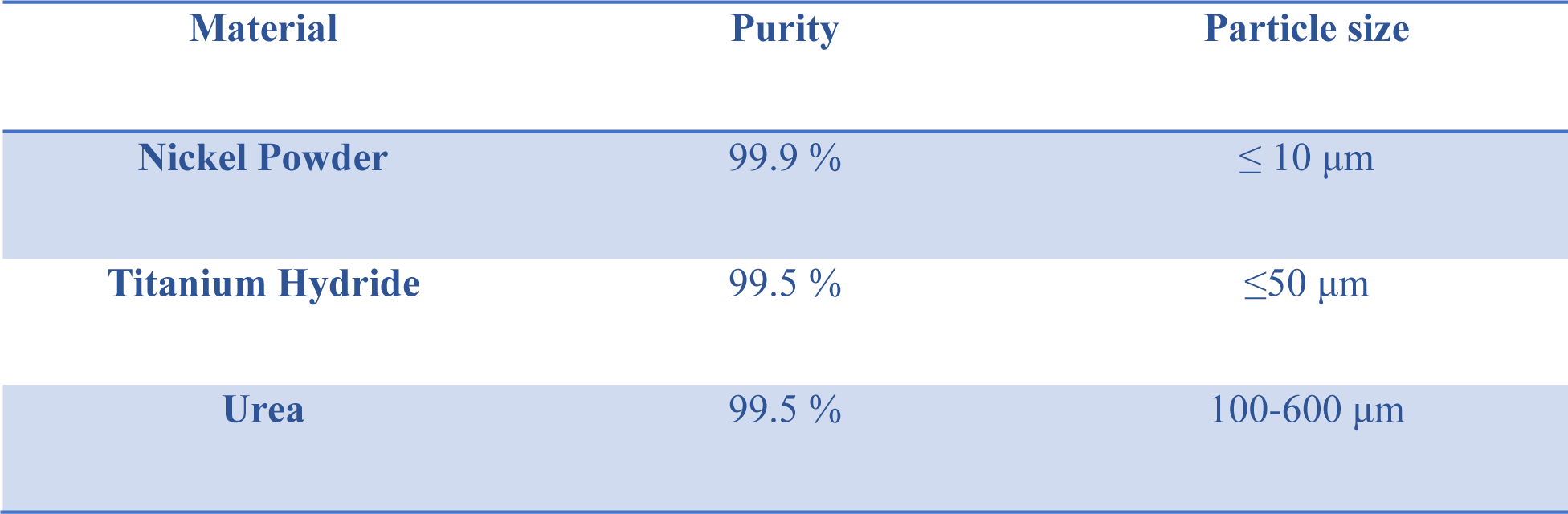
Provided from Merck the urea, titanium hydride, and nickel powder.

| Material | Purity | Particle size |
| --- | --- | --- |
| Nickel Powder | 99.9 % | $\leq 10 \mu\text{m}$ |
| Titanium Hydride | 99.5 % | $\leq 50 \mu\text{m}$ |
| Urea | 99.5 % | 100-600 $\mu\text{m}$ |

### 2.2 Surface Analysis

The scanning electron microscope (SERON TECHNOLOGY, AIS2100, Korea) was used to analyze the surface morphology of the polished nitinol sample. In the processing method, Urea was added as a porosity maker to oblige the sample to be more compatible with the body in terms of stress shielding and cell adhesion. As every sample exhibits non-uniform surface conditions that are defined by features resembling dimples, the polishing procedure was applied. Then the sample undergoes a gold coating process using a technology that applies a 20 nm thick layer of gold by Physical Vapor Technology (PVD). The Nitinol specimen was examined using Digimizer 6 software to evaluate the pore size and its competence with the desired employment.

### 2.3 The AFM test

In this phase, an experimental evaluation was conducted on WB and WOB samples to assess the surface roughness. The experiment was conducted using an Atomic Force Microscopy (AFM) apparatus manufactured by JPK business, specifically the Nanowizard 3 model. The first picture was captured using the non-contact imaging mode, resulting in a size of 10 × 10 micrometers and a resolution of 128 dots per inch (dpi). The subsequent region was chosen inside the first picture, measuring 1 × 1 μm, and was then captured at a resolution of 256 dots per inch (dpi). The analysis was performed with JPK Data Processing (version 5.0.96) and two-dimensional, three-dimensional images and their histogram frequency chart were presented in Figure 3. The average surface roughness (Ra), the root means square roughness (RMS), and peak-to-valley roughness (Rt) are obtained by the software as indicated in Table 2.

**Figure 3a.**
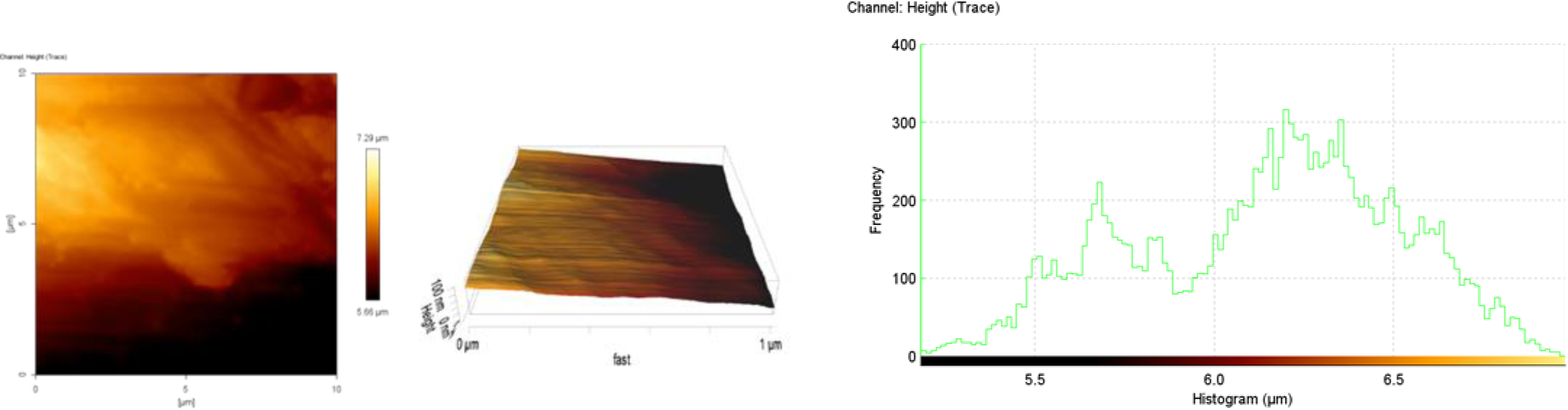
2D and 3D AFM imaging and height histogram of NiTi before braiding (WOB).

**Figure 3b.**
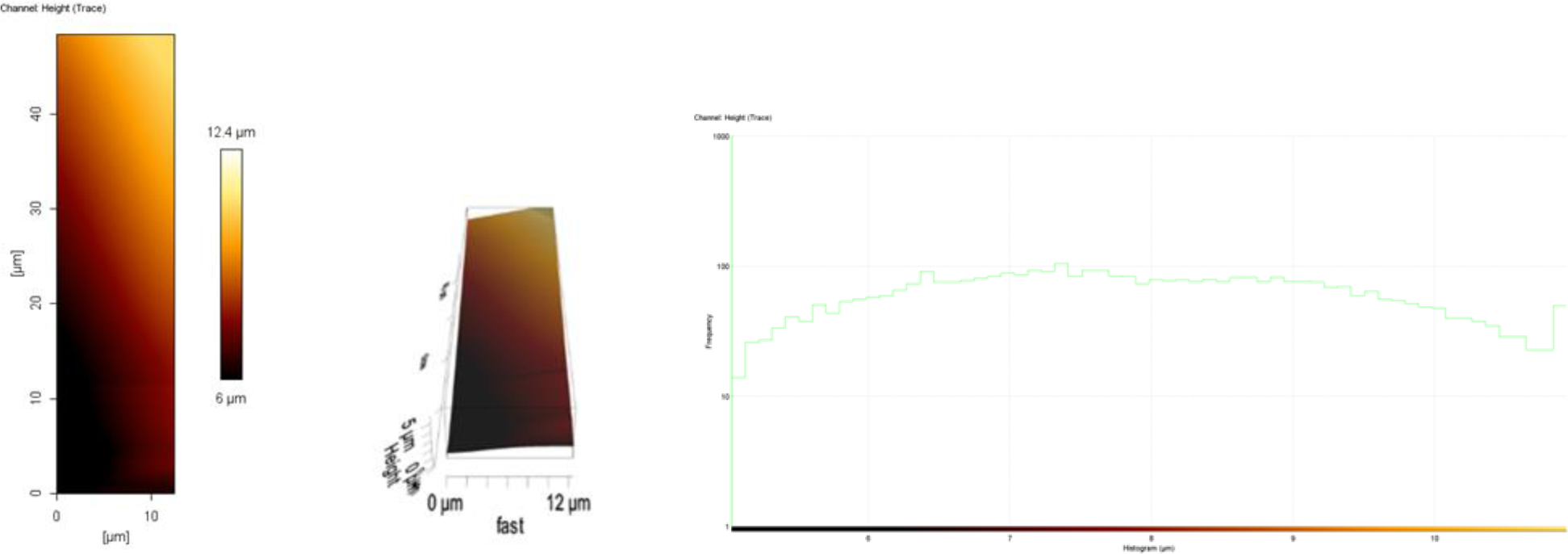
2D and 3D AFM imaging and height histogram of NiTi after braiding (WB).

**Table 2.** NiTi roughness information WB and WOB.

| Roughness Information | A) Height (WOB) | B) Height (WB) |
| --- | --- | --- |
| Average Value | 6.144 $\mu\text{m}$ | 7.907 $\mu\text{m}$ |
| Average Roughness Ra | 267.5 nm | 975.1 nm |
| RMS Roughness Rq | 370.6 nm | 1.433 $\mu\text{m}$ |
| Peak-to-Valley Roughness Rt | 1.782 $\mu\text{m}$ | 5.803 $\mu\text{m}$ |

### 2.4 Investigation of corrosion by electrochemical impedance testing

The electrochemical impedance test involves introducing alternating current into the system and receiving impedance from the system in two forms: real and imaginary impedance. It is possible to examine the effective parameters for corrosion of the system by fitting the findings on the circuit of electrochemical equivalents by charting this information in terms of each other in the following distinct ways. To evaluate the resistive behavior and electrochemical activity during implant corrosion after placement in the body, samples WOB and WB from electrochemical impedance spectroscopy (EIS), a tool for understanding deterioration, were used. The Ivium device (Potentiostat-Galvanostst VERTEX-One) performed this test on a surface of 1cm2 with a frequency range of 10 mHz to 100 kHz and 10 mV amplitude sine wave.

### 2.5 Cell adhesion test

L929 cells (NCBI C161) from the Iranian Pasteur Institute’s cell collection were used in this study. The cells were frozen, then they were moved to a flask filled with RPMI media containing 10% FBS. The flask was then kept in an incubator set at 37°C, 90% humidity, and 5% carbon dioxide concentration. Note that the culture media is replaced every three to four days. First, alcohol was used to disinfect the samples. Sterilized samples (WOB and WB) were inserted into each well of the sterile 24-well plate to assess cell adherence. Following the addition of 2 x 104 cells in a volume of 60 microliters to each sample, the samples were incubated for 4–5 hours in a Memlert incubator. Following the time for the cells to adhere, a particular volume of culture media containing 10% FBS was added to each well. The samples’ growing medium was removed after 24 hours, and they were then cleaned with phosphate-buffered saline (PBS) for 30 seconds before being fixed with 3.5% glutaraldehyde. A particular amount of fixative was applied to each sample, and after being refrigerated for two hours, the fixative was removed. The samples were then treated with deionized water and alcohol washes at 50%, 60%, 70%, 80%, and 96%. Cell adhesion to the samples was checked with a scanning electron microscope (SERON TECHNOLOGY) model AIS2100 at Amirkabir University of Technology at different scales and magnifications.

### 2.6 MTT Assay

Due to the findings of research indicating the robust biocompatibility of nitinol, which is a crucial criterion for implants inside the human body, an in vitro test for further examination, was applied. In this experimental setup, the sample is brought into contact with MG63 cells that obtained from the cell bank of Pasteur Institute of Iran, and afterward, the viability of the cells will be evaluated by the use of an MTT kit. The cells are cultivated in culture flasks inside an incubator set at a temperature of 37 °C and an atmosphere of 5% carbon dioxide, maintaining a humidity level of 85%. The culture media used is RPMI-1640, which has 50 units of penicillin and 50 g of streptomycin per milliliter. 10% of fetal calf serum (FCS) is also added to the culture medium. Following the creation of the cell layer, which takes three to four days, the cells are extracted from the flask’s surface using the trypsin enzyme (0.25%), and a solution containing four times the number of cells per millimeter is ready for use. Conversely, WB and WOB samples are oven-sterilized (dry heat). The samples were placed in a 12-house container (each sample in a well) and a well without a sample is considered to be a control. After placing the samples in each of the wells, 2 ml of cell suspension is poured and placed in the incubator. Then MTT solution is added to the wells and they are incubated again. A microtiter reader (Merck, Germany) is used to read the content of the corresponding plate containers after 200 microliters of isopropanol solution (Merck, Germany) is added to the corresponding wells following 3 to 5 hours of incubation at 37 °C. The corresponding plate containers are then placed on the shaker for 10 to 15 minutes. A solution of MTT containing 5 mg/ml is made by dissolving 50 mg of MTT powder in 10 mg of 150 x 10-3 M PBS. When using it during the test, it is diluted up to 10 times with PBS to make a solution of 0.5 mg/ml MTT. The MTT method is usually used to check the survival of cells. In this method, tetrazolium yellow salt is employed. When the cells absorb it, a color change occurs and this salt turns purple. The reason for the color change is the formation of insoluble crystals. This color change can be detected by spectroscopic methods. For each cell, there is a linear relationship between the number of living cells and the amount of light absorption [25]. It should be noted that this test was done on five periods, namely the first, third, seventh, fourteenth, and twenty-first days on the WB and WOB samples to compare their effect on biocompatibility.

## 3. Result and Discussion

### 3.1 Morphology study

Figure 4 illustrates the SEM photographs of the materials. Table 3 provides details on the parameters of the pores, such as their length, perimeter, and area. The significance of pore size concerning cell development is well-established. When the pore size is small, cells have limitations in their ability to migrate towards the central region of a structure. This, in turn, hampers the diffusion of nutrients and the elimination of waste materials. On the other hand, if the pores exhibit excessive size, there is a resultant decrease in the specific surface area, hence imposing constraints on the attachment of cells. By choosing two pores in different magnifications and calculating with Digimizer 6 software, the average pore size observed in the sample is around 310 micrometers that indicated by the area of 48636.3142 square micrometers that by comparing this size with the size of pores in the bone [26] it can be demonstrated that the congruity between these two amounts is the preference of an orthopedic implant to have osseointegration and elimination of stress shielding. According to study [27], the best pore size for bone growth is 300-800 micrometers, which specifies the manufactured sample’s suitability as a bone implant. This particular pore size has been shown to facilitate the optimal proliferation of bone cells and their adhesion [28, 29]. As a fact, cell adhesion plays a role in various natural processes, including maintaining tissue structure, healing wounds, and integrating biological materials into tissues. The biocompatibility of biomaterials is intricately related to the interaction between cells and the surface of the material, especially in terms of cell adhesion. Surface properties of materials, including their topography, are important factors in determining the adhesion of osteoblasts to biomaterials. This feature also affects bone cells [30], which can be seen in biological SEM images (Fig. 7). While there may be different forms in the behavior of diverse cells, it can be generally said that an appropriate sample will be available to support and facilitate bone cell activities in which the result of the applied biological tests also confirms the established pores in the samples coincide with its application as an implant in hard tissue and caused cell mobilization. So, adding the Urea as a porosity maker does its work in the desired way for enhancing biological properties.

**Figure 4.**
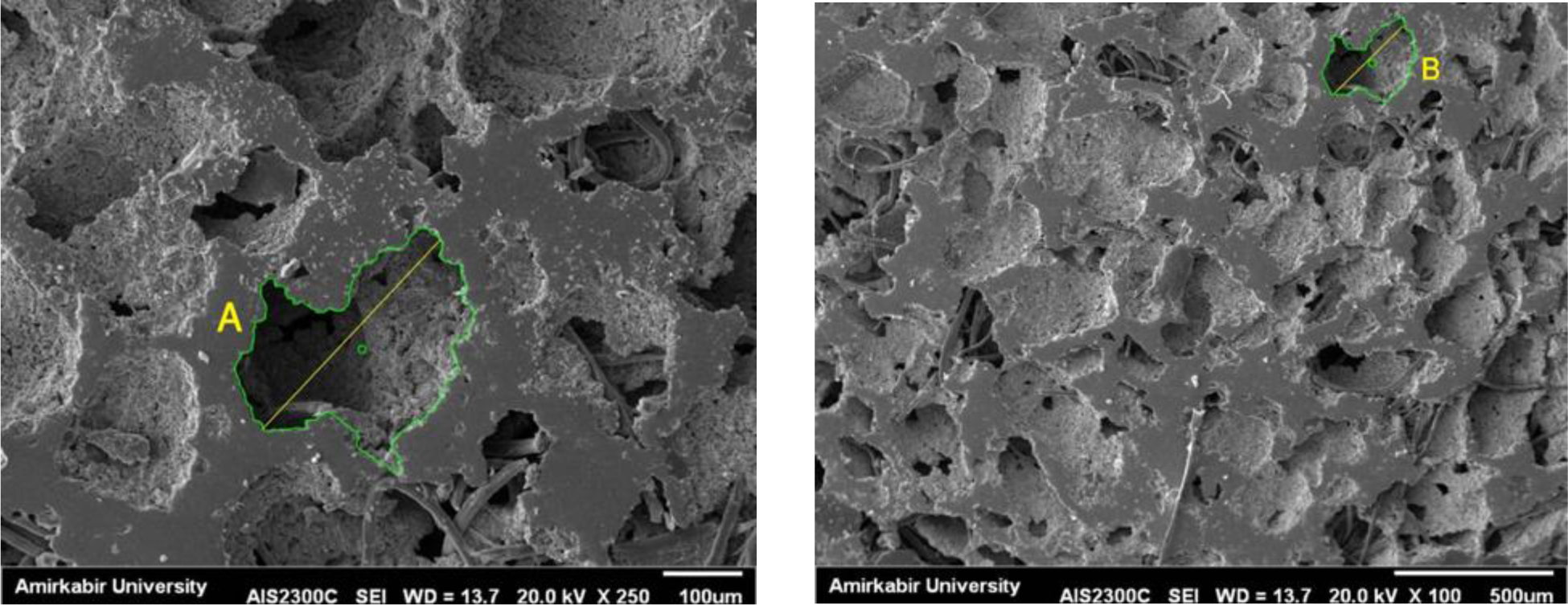
the SEM images taken at two different magnifications (A: 100 µm and B: 500 µm).

**Table 3.** Specifics of the pore parameters—including length, area, and perimeter.

|  | Length | Perimeter | Area |
| --- | --- | --- | --- |
| **A (100 $\mu\text{m}$)** | 321.629 $\mu\text{m}$ | 1150.7824 $\mu\text{m}$ | 49382.4188 $\mu\text{m}^2$ |
| **B (500 $\mu\text{m}$)** | 300.126 $\mu\text{m}$ | 1006.9116 $\mu\text{m}$ | 47890.2095 $\mu\text{m}^2$ |
| **Mean** | 310.8775 $\mu\text{m}$ | 1078.847 $\mu\text{m}$ | 48636.3142 $\mu\text{m}^2$ |

### 3.2. AFM assay

The information about roughness is succinctly presented in Table 2. The average roughness for sample WB and WOB is 267.5 nm and 975.1 nm respectively (p < 0.05). By comparing these numbers, a significant increase in the roughness value was observed by appending the braid to the cylinder. Mainly by comparing the table amounts, the surface roughness becomes greater in WB. The AFM topography images and histogram of WOB and WB samples are assessed (Fig. 3a and 3b) to evaluate the nanodomain dimensions. The objective of this study is to optimize protein adsorption and cell adhesion, making this issue of utmost importance that the rise in roughness results in better cell adhesion (SEM images in Figure 7 confirm). To modify the rate of coagulability and enhance the prompt synthesis of cells and tissues, it is necessary to make modest adjustments to the porosity and roughness. As roughness is a notable factor needed for the manufacturing of metal implants that affects bone integration, by investigating the results achieved from this test and SEM analysis, it can be stated that the feature in the samples with braid can be suitable for implementation in the bone area.

### 3.3. Corrosion test results of electrochemical impedance spectroscopy

As shown in Fig. 5a, the Nyquist curves (Z’ in terms of -Z”) related to samples WB and WOB, the frequency increases from the right side of the graph to the left side of the curve (clockwise), so the point located at the far left of the graph has the highest frequency and the extreme point on the right has the lowest frequency. In these curves, the increase in diameter indicates the expansion in the corrosion resistance of the system [31]. Therefore, it is clear from the shape of the Nyquist curves that the diameter of the curve in the WB sample is higher than WOB. The plots of bode-impedance and bode-phase angle related to these samples are shown in Fig. 5b. In the impedance bode curves, the impedance at the lowest frequency can represent the resistance of the entire system against corrosion [31]. Therefore, it is clear from Fig. 5b (A) that the impedance at the lowest frequency of the WB sample was higher than WOB. In addition, WOB had the lowest impedance at the lowest frequency. By analyzing the parameter values obtained from this test (Table 4), it is clear that WB has a higher load transfer resistance than WOB. This higher load transfer resistance means higher resistance of this sample against corrosion. So, it can be concluded that the presence of the braid has increased the corrosion resistance for instance R_ct_ in WOB and WB is 6352 and 7366 ohm.cm^2^, respectively. The exact value of the resistance of these systems should be determined by modeling these results by the electrochemical equivalent circuit (Fig. 6). Modeling of measured samples with electrochemical equivalent circuit was done by ZsimpWin software and the results are shown in Fig. 5a. and 5b. As can be seen, the modeling has been able to match the Nyquist and Bode diagrams well, which indicates the reliability of the modeling results. To discover the shape of the electrochemical equivalent circuit, it is necessary to find the number of time constants of the system which means the number of parallel capacitors/resistors in the electrochemical equivalent circuit. This circuit includes two resistors (the solution resistor and the charge transfer resistor from the left, respectively) and a constant phase element (related to the double layer). In this equivalent circuit, because the plates of the double layer created between the surface of the electrode and the electrolyte are not exactly parallel to each other and the plate of the electrode and the cover are not completely smooth, a phase constant element is used instead of an ideal capacitor.

**Figure 5a.**
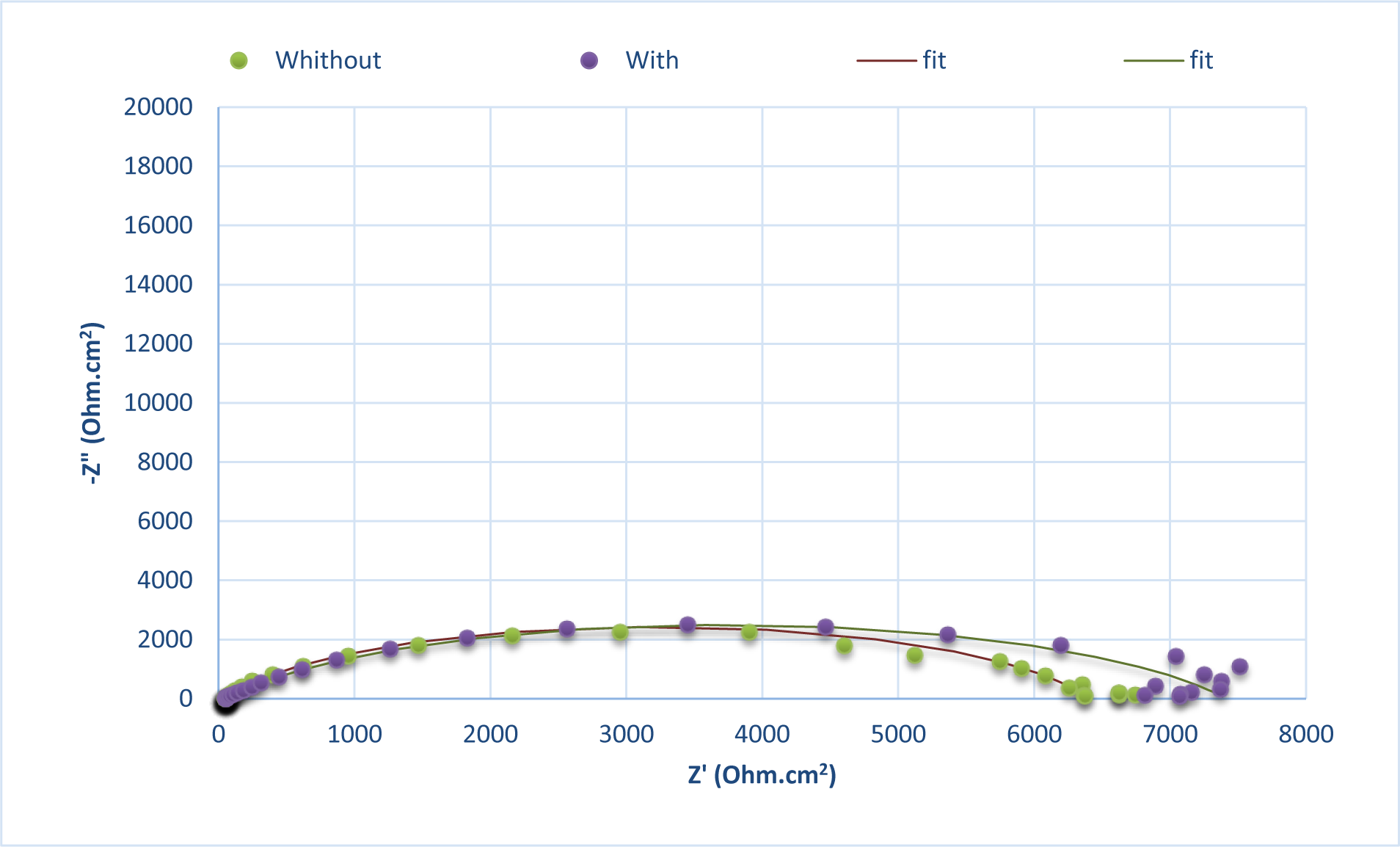
Nyquist curves of the studied samples in addition to the fit results of the data obtained from the EIS test on the appropriate electrochemical equivalent circuit (the points are the results of the test and the lines are the results of the fit).

**Figure 5b.**
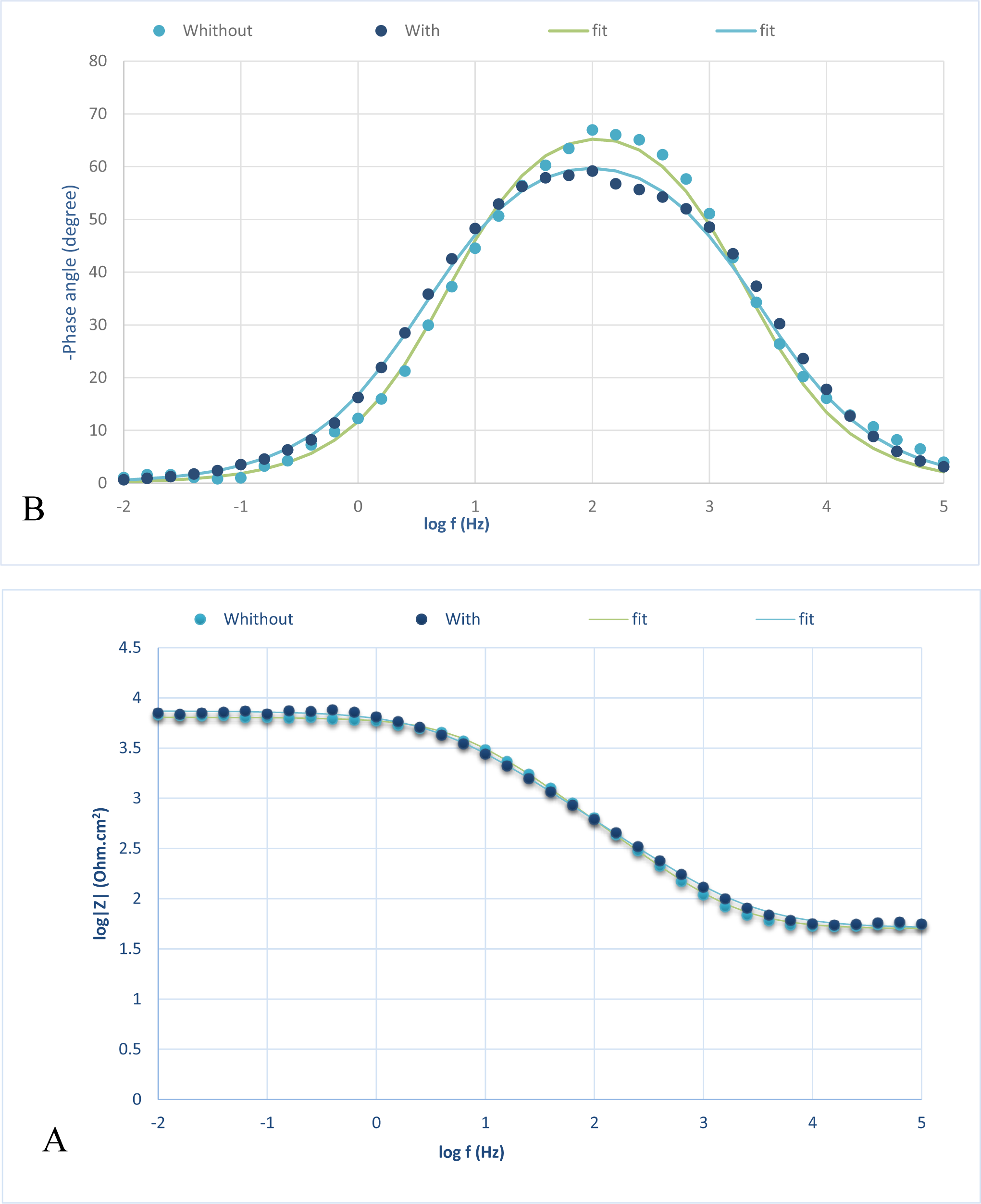
The curves of (A) bode-impedance modulus and (B) bode-phase angle of the samples (the points are the results of the test and the lines are the results of the fit).

**Figure 6.**
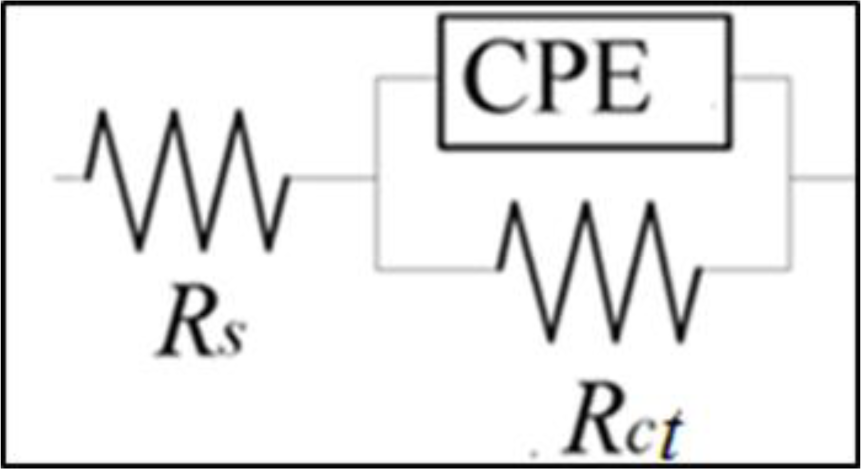
The electrochemical equivalent circuit used to model the results of the electrochemical impedance test.

**Table 4.** The results of modeling the impedance test results on the electrochemical equivalent circuit.

| Sample | $R_s$ (ohm.cm<sup>2</sup>) | $Y_{0dl}$ (S.sec<sup>n</sup> .cm<sup>-2</sup>) | n | $R_{ct}$ (ohm.cm<sup>2</sup>) |
| --- | --- | --- | --- | --- |
| without braided cover | 49.91 | 7.82E-06 | 0.83 | 6352 |
| with braided cover | 50.84 | 1.25E-05 | 0.76 | 7366 |
| NiTi wire | 48.68 | 5.68E-06 | 0.81 | 14560 |

These factors manifest themselves in the impedance formula. The impedance of a capacitor is determined by (*Z*) is calculated using the equation:

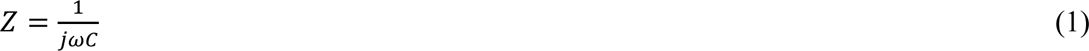

Similarly, the value for CPE is determined by (*Z*) formula is calculated using the following formula:

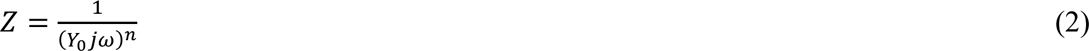

(*C*: capacitor capacity, *ω*: the phase angle, *Y*_0_: the admittance, *j*: the imaginary term 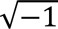).

The distinction between these two digits is only based on the variable “*n*”, which represents a numerical value ranging from zero to one. An ideal resistor is represented by a value of zero, whereas an ideal capacitor is represented by a value of one.

### 3.4. Cell adhesion test

In the SEM images taken in Figure 7, the adhesion of the cells is visible, in the sample WB (b), this adhesion can be seen compared to the sample WOB (a) at the same scale and magnification, and there is an increase. The braid has provided more suitable conditions for the proliferation of cells and their growth. In two images (c) and (d), we can see the sample WOB and WB, respectively, with the same scale but in different magnifications. According to the results of surface analysis taken by SEM, the pore size of the sample agrees with cell desire for adhesion that has been confirmed with these images. It can be inferred that the use of urea with a porosity size ranging from 100-600 μm leads to the formation of pores with a size of 300 μm, which aligns with the desired porosity for bone tissue and is consistent with the porosity of bones in the body.

**Figure 7.**
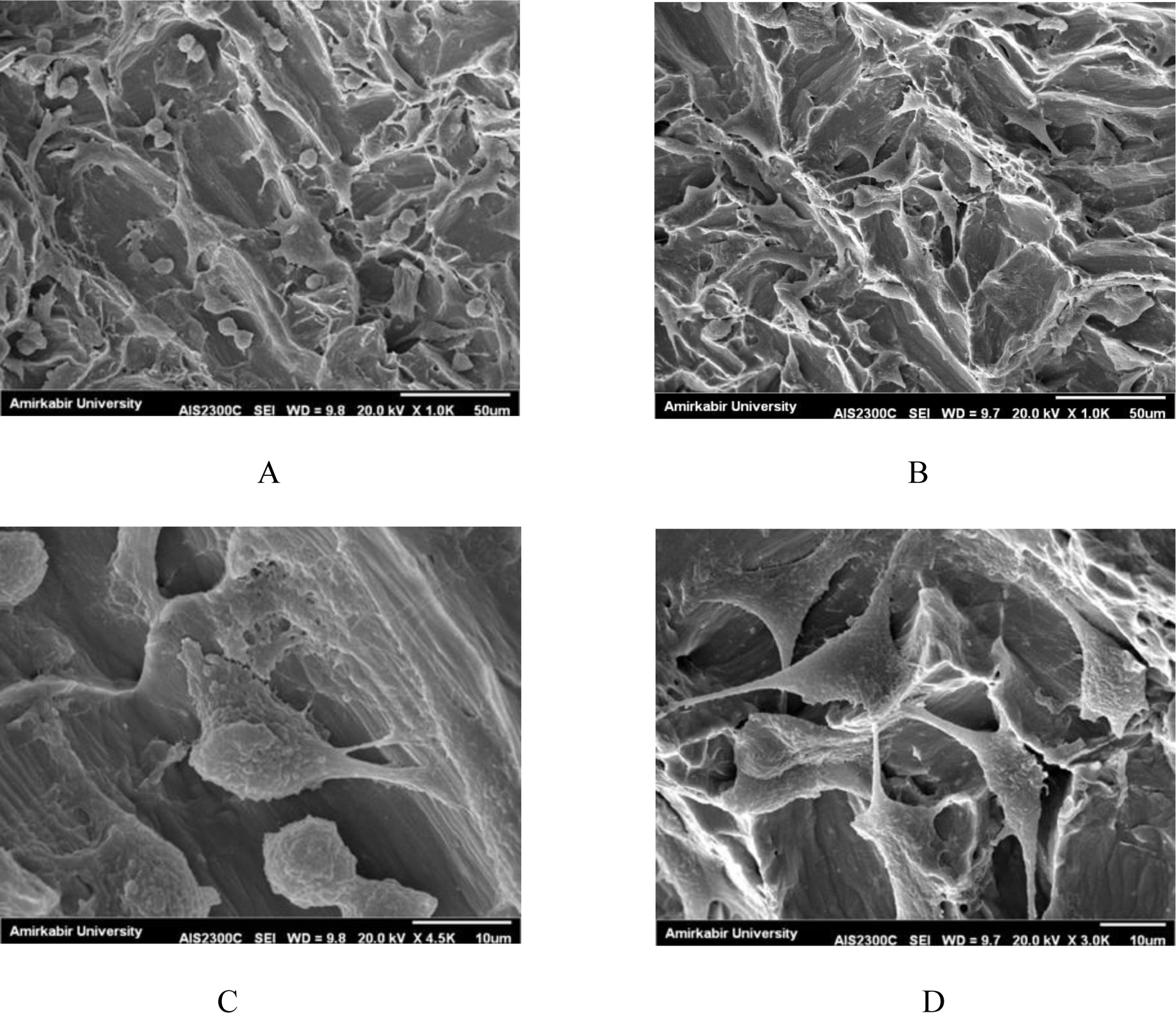
The SEM images depict the fibroblast cell culture. Image A (WOB) and image B (WB) with a magnification of 50 µm, while image C (WOB) and image D (WB) with a magnification of 10 µm. All photos are shown in the same scale.

### 3.5. MTT cell test result

MTT test was performed on both samples. The graph in Figure 8. shows the percentage of cell survival according to the time of one, three, seven, fourteen and twenty-one days. In the WOB sample, within three days after the initial cultivation of cells on the sample, the percentage of cell survival has reached from 33% to 48.3%, and in the other sample, this percentage has increased from 35 to 47. This means that on the first day, the percentage of survival in the sample with a cover was higher (35% versus 33%), but after 72 hours, the percentage of survival was the same (around 48%). In the following, it seems that after 7 and 14 days, both samples have reached a high percentage of survival rate and are very close to each other, and after 21 days, the peak survival percentage is 95.1% and 93.9% in WOB and WB, respectively. Thus, in addition to the nitinol sample’s low toxicity and high cell survival rate, it can be inferred that the inclusion of the braid did not result in cell death and that the cell survival rate was ultimately equal to that of the sample without the braid. In other words, the growth and proliferation of cells in both samples are suitable, which indicates that the cells did not suffer from toxicity and their viability was not overshadowed because if the sample had a high level of toxicity, apart from cell death, the cells would not have the ability to grow and reproduce. The result is in alignment with the objective of the study to develop an implant that possesses both biological and mechanical properties similar to bone, specifically tailored for this purpose.

**Figure 8.**
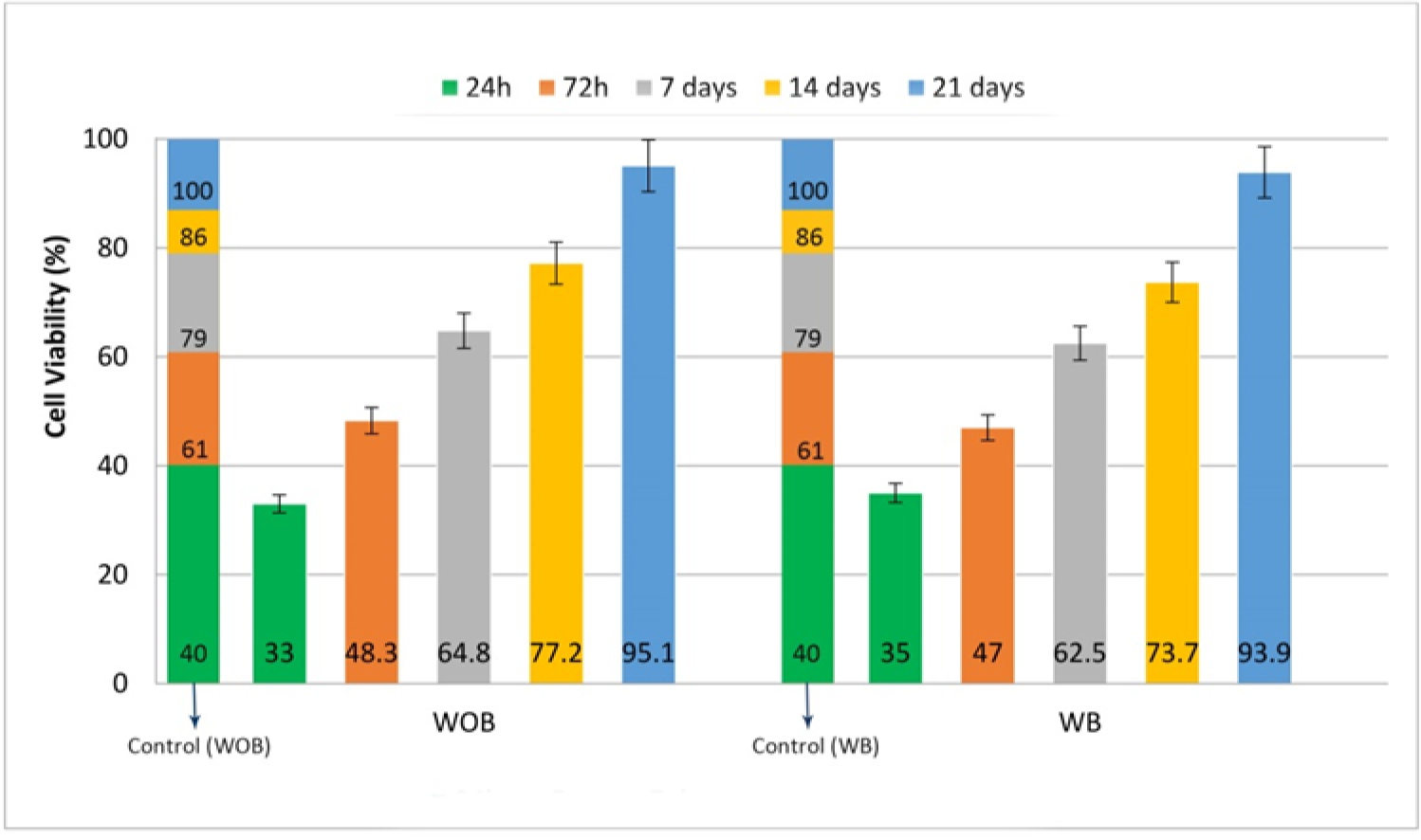
Cell viability evaluation using the MTT test at 1, 3, 7, 14 and 21 days after MG 63 cell culture (p < 0.05).

## 4. Conclusion

In this study, we developed and evaluated a novel Nitinol lumbar vertebral implant using a two-step sintering and braiding process. Our findings demonstrate that the incorporation of a braided structure significantly enhances both the mechanical and biological properties of the implant.

Surface morphology and roughness analyses revealed that the braided samples exhibited a pore size of approximately 310 micrometers, as observed through SEM analysis, and an average roughness of 975.1 nm, as measured by AFM. These features are conducive to bone cell adhesion and growth, in contrast to the non-braided samples, which had a roughness of 267.5 nm.

Corrosion resistance was assessed using Electrochemical Impedance Spectroscopy, which showed that the braided samples had a higher charge transfer resistance (R_ct_ = 7366 ohm.cm²) compared to the non-braided samples (R_ct_ = 6352 ohm.cm²), indicating superior corrosion resistance.

Biocompatibility tests, including cell adhesion and MTT assays, confirmed the non-toxicity and biocompatibility of the braided samples, which exhibited a cell viability rate of 95.1% over 21 days.

These results suggest that braided Nitinol implants offer a promising approach for lumbar vertebral implants, combining enhanced mechanical properties with excellent biocompatibility. The braiding technique significantly improves the overall performance of the implants, potentially leading to better outcomes in spinal surgeries. Future research should focus on optimizing the braiding process and conducting long-term in vivo evaluations to further validate these findings and enhance clinical applicability.

## AUTHOR INFORMATION

### Author Contributions

Conception or design of the work, Writing - original draft, and Writing - review & editing. ‡ Mahdis Parsafar

Data collection, Analysis and interpretation, Writing - original draft, and Writing - review & editing ‡ Alireza Tehranian

Data collection, Drafting the article, Writing - original draft, and Writing - review & editing. ‡ Ramin Ardalani

## ABBREVIATIONS

LBP: Low Back Pain
BTE: Bone Tissue Engineering
WOB: With Out Braid
WB: With Braid

